# Harvest and Culture of Porcine Adipose-Derived Mesenchymal Stem Cells for Autologous Transplantation

**DOI:** 10.1101/2020.06.25.169367

**Authors:** Ayushman Sharma, Allan B. Dietz

**Affiliations:** Division of Experimental Pathology and Laboratory Medicine (Drs Sharma and Dietz), Division of Transfusion Medicine (Dr Dietz), and Department of Immunology (Dr Dietz), Mayo Clinic, Rochester, Minnesota

**Keywords:** ADMSCs, MSC, pADMSC, porcine fat stem cells

## Abstract

**Importance:** Transplantation of adipose-derived (mesenchymal) stem cells (ADSCs) are currently under investigation for numerous novel regenerative cell therapies of the head, neck and periphery. Critical to the development of these techniques is the availability of large-animal models that can be used to test the safety and efficacy of these approaches in a manner that provides source material (in this case MSC) analogous to those developed in humans.

**Objective:** To describe the surgical technique and laboratory procedures for harvesting and isolating porcine ADSCs that are functionally equivalent to human ADSCs without sacrificing the donor animal.

**Methods:** The reagents and methods used in the porcine model described were purposefully focused to be able to be sufficiently analogous to those used in humans such that data developed using these techniques should support the use of porcine models for regulatory submissions.

**Results:** We describe a method and confirm the activity of functionally analogous adipose derived porcine MSC. Two conditions were critical to move gain analogous performance: the cells needed to be incubated at porcine body temperature (39°C) and the cells were more sensitive to initial plating densities with plating densities of 20,000 cells/cm^2^ being optimal.

**Discussion:** This approach will allow reproducible and predictable use of an autologous large-animal model for testing AMDSC therapies.

## Introduction

Autologous adipose-derived stem cells (AMDSCs; also termed *mesenchymal stem cells* [MSCs]) are easily obtained, multipotent, and versatile cells that have shown remarkable healing properties in varied clinical settings [1, 2], but the development of effective clinical therapies requires rigorous in vitro and preclinical in vivo studies. Domestic swine are the ideal animal model for many human disease states because of their anatomic and physiologic similarities to humans. Hence, isolating, harvesting, and testing of porcine ADMSCs (pADMSCs) are crucial techniques when evaluating the safety and efficacy of new cell therapies.

Several papers have briefly discussed porcine fat isolation and characterization but only as part of an in vivo experiment or in experiments without survival surgeries. Many of these studies have used culture conditions identical to those used for humans [3-13]. Here, we provide specific surgical techniques for porcine fat harvest as a survival surgery and describe pADMSC isolation and culture optimization for autologous transplantation, including culture conditions that produce porcine MSCs that are analogous to those cultured from humans, with similar proliferation and differentiation capacities.

## Materials and Methods

This study was approved by the Mayo Clinic Institutional Animal Care and Use Committee (protocol A00002304-17). Institutional policies for animal handling were followed.

### Adipose Tissue Harvest

These techniques were developed based on their successful use in humans [1, 14, 15]. Healthy domestic white pigs (*Sus scrofa domesticus*) without any previous surgical procedures, weighing from 40 to 80 kg, were anesthetized with intramuscular tiletamine-zolazepam (6 mg/kg) and xylazine (2 mg/kg), followed by intubation and anesthesia maintenance with isoflurane. Animals were then positioned either in the dorsal recumbent position or ventral recumbent position (Figure 1A & Fat Harvest Video). The dorsal recumbent position was used when harvest was planned from the upper chest area, whereas the ventral recumbent position was used when harvesting fat from the back. For domestic white pigs, these 2 areas are collectively termed the *fat back* [16]. Unlike human fat, porcine fat is tightly adherent to the overlying dermis and muscular fascia below. For animals weighing less than 40 kg, the back is the ideal position to harvest adipose tissue because the chest area has a much thinner layer of subcutaneous fat.

**Figure 1.**
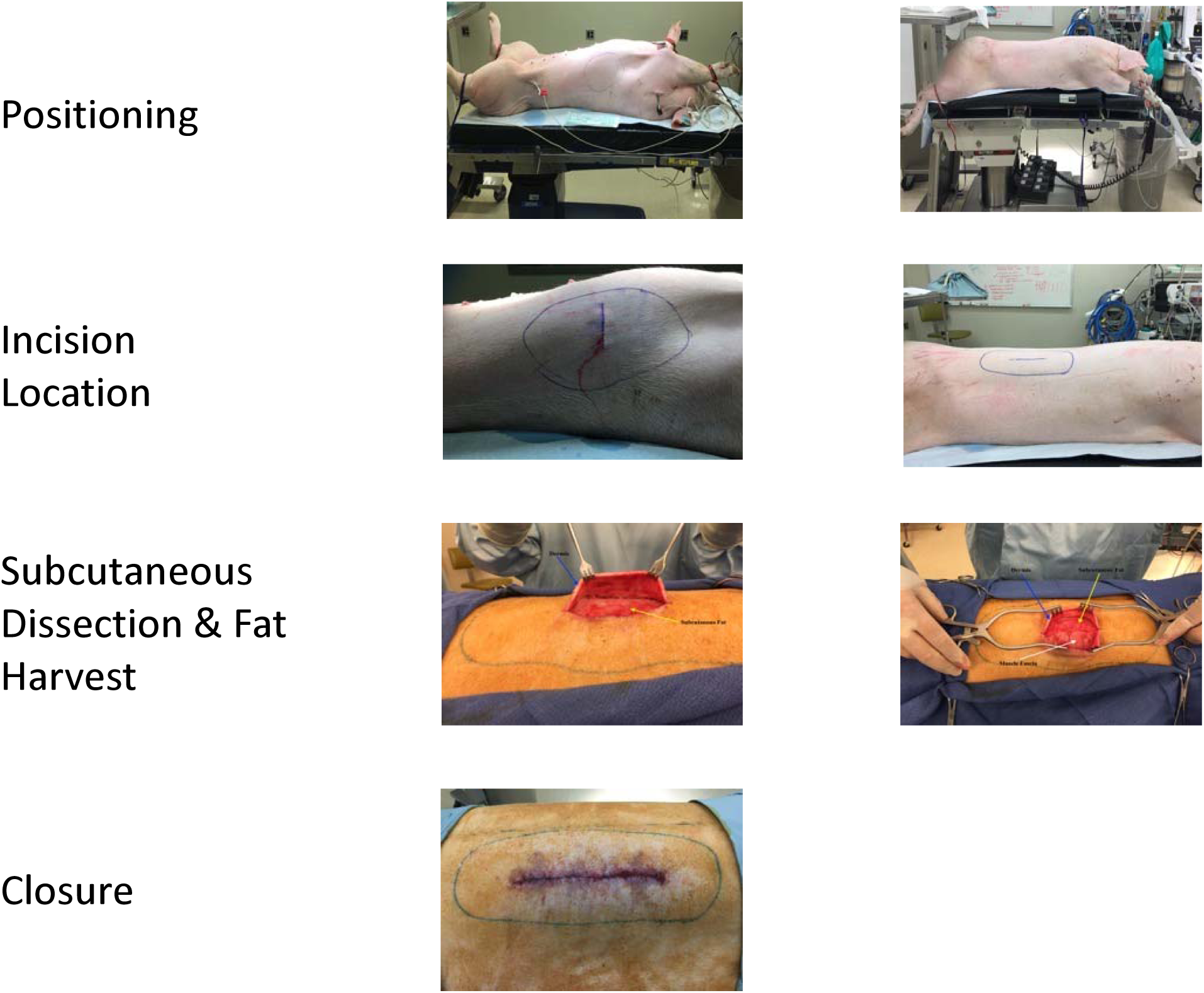
Surgical Procedure for Porcine Adipose Tissue Harvest. Subcutaneous fat from adult pigs can be harvested from the upper chest or the back. Harvest from the chest requires positioning the animal in a dorsal recumbent position, whereas harvesting from the back requires a ventral recumbent position. The incision should be made to expose subcutaneous fat, and dissecting scissors should be used to separate the tightly adherent fat from the overlying dermis while leaving the muscular fascia intact. Skin should be closed with 2 layers of sutures and skin adhesive to prevent contamination. A, Positioning: dorsal recumbent (left) and ventral recumbent (right). B, Incision location. C, Subcutaneous dissection and fat harvest. D, Closure.

After positioning, an 8-cm incision was marked with a surgical marking pen (Figure 1B). The skin was cleaned with antibacterial solutions and draped in a sterile fashion to avoid fecal contamination of the sample. A no. 20 scalpel was used to make an incision through the dermis to expose underlying subcutaneous fat. Skin rakes or skin hooks were then used to retract the skin while sharp dissecting scissors were used to separate the fat from the subcutaneous tissue and underlying muscular fascia (Figure 1C). Depending on the weight of the pig, we collected 7.5 to 15 g of fat from each pig. After harvest, the tissue was placed in sterile saline or lactated Ringer solution for transport to the laboratory for processing.

The incision was closed in 3 layers: a deep dermal layer, a cutaneous layer, and a cyanoacrylate skin adhesive (Dermabond; Ethicon Inc; Somerville, New Jersey) (Figure 1D). The wound was then covered by antimicrobial incise drapes (Ioban; 3M Medical; St Paul, Minnesota) to further reduce chance of fecal contamination. Animals were kept in separate kennels for 2 weeks while the wounds healed. Other than keeping the antimicrobial drapes in place for 2 weeks, the pigs had no special wound care needs. No adverse events (eg, wound dehiscence, seroma, hematoma, infection) were associated with the harvest procedure.

### Tissue Processing and ADMSC Isolation

All tissue processing was performed in a clean and sterile hood with sterile equipment to reduce risk of culture failure from contamination. (Fat Processing Video) Phosphate-buffered saline (PBS; 100 mL) was mixed with 75 mg (±5 mg) of type A collagenase (Worthington Biochemical Corporation; Lakewood, New Jersey), filtered through a 0.2-μm filter, and placed in a 500-mL flask.

Porcine adipose tissue (up to 10 g) was cleansed by vigorous vortexing (10- 30 seconds) in PBS and centrifugation (5 minutes at 260×*g*). The fat sample was minced as completely as possible with dissecting scissors and placed in the flask containing the collagenase A solution. The mixture was incubated at 37°C for 60 minutes, with intermittent swirling by hand every 5 to 7 minutes.

At the end of the incubation, the collagenase was neutralized with an equal volume of culture media, which consisted of advanced minimum essential media (Gibco; Camarillo, California), 5% human platelet lysate (PLTMax; Mill Creek LifeSciences; Rochester, Minnesota), 1% GlutaMax (ThermoFisher Scientific; Waltham, Massachusetts), 1% penicillin/streptomycin (ThermoFisher Scientific), and heparin sulfate (App Pharmaceuticals; Lake Zurich, Illinois). This mixture was transferred into 50-mL conical tubes and centrifuged (10 minutes at 1,125×*g*, room temperature). The supernatant (including the floating adipose layer) was aspirated, and the pellet was resuspended in 10 mL of culture media.

Resuspended pellets were combined and filtered (70-μm Falcon cell strainers; Corning Enterprises; Corning, New York). Filtered cells were collected and resuspended as described above, and cells were refiltered (40-μm Falcon cell strainers; Corning Enterprises). Upon final collection (centrifugation for 10 minutes at 1,125×*g*), the pellet was resuspended in 30 mL of culture media. Resuspended cells were plated into T-75 flasks (5 g of processed fat per T-75 flask) and incubated at 39°C (normal porcine body temperature) with 5% CO_2_.

### Growth Kinetics of pADMSCs

Growth rates of human and porcine ADMSCs were compared by using the IncuCyte Zoom cell imaging system (Essen Bioscience; Ann Arbor, Michigan). Passage 4 MSCs from 4 human patients and 4 pigs (Table 1) were plated in 6-well plates at the following concentrations: 20,000 cells/cm^2^ (high density), 10,000 cells/cm^2^ (medium density), and 5,000 cells/cm^2^ (low density). Standard culture media were used for all conditions [17]. Human cells were incubated at 37°C in the IncuCyte Zoom system, and confluence was measured every 12 hours with phase-contrast imaging. Porcine cells were treated identically but incubated at either 37°C or 39°C.

**Table 1:** Age and Gender of Human and Porcine Donors.

| **Donor Designation** | **Age, y/m** | **Sex** |
| --- | --- | --- |
| **H-1** | **58y** | **Female** |
| **H_2** | **61y** | **Female** |
| **H_3** | **60y** | **Female** |
| **H_4** | **37y** | **Female** |
| **P_1** | **5m** | **Male** |
| **P_2** | **4m** | **Female** |
| **P_3** | **3m** | **Female** |
| **P_4** | **5m** | **Female** |

### Characterization of pADMSCs

The multipotent potential of human ADMSCs (hADMSCs) was verified by using their capacity to differentiate into adipocytes, osteocytes, and chondrocytes [18]. Although porcine and human cells do share homology, few antibodies have been verified to have cross-species reactivity and reproduce comparable flow cytometry results. The multipotent potential of pADMSCs was likewise evaluated by using trilineage differentiation.

Differentiation procedures for adipogenesis and osteogenesis were performed in triplicate, following manufacturer protocols (StemPro Adipogenesis & Osteogenesis Kits; Gibco), with the following exceptions to optimize for pADMSC differentiation [19, 20]:

a. Initial plating density for both differentiation procedures was 20,000 pADMSCs/cm^2^
b. Cells reached 70% confluence before differentiation media was added
c. Incubation temperature for differentiation was 39°C for 1 set of pADMSC cultures and 37°C for all others
d. Adipogenesis differentiation was carried out for 14 days, and osteogenesis differentiation was carried out for 18 days.

Oil-Red-O [21] and Alizarin red [22] staining were performed according to manufacturer protocols.

Differentiation procedures for chondrogenesis were followed in triplicate according to manufacturer protocols (StemPro Chondrogenesis Kit; Gibco), with the following exceptions to optimize for pADMSC differentiation [23]:

a. Micromass cultures were created using 15 μL of pADMSCs suspended at 1×10^7^ cells/mL vs 5 μL, as suggested by the manufacturer
b. Incubation temperature for differentiation was 39°C for 1 set of pADMSC cultures and 37°C for all others.

Alcian blue staining was performed by using components from the Alcian blue kit (Vector Laboratories; Burlingame, California) [24]. Briefly, micromass cultures were fixed with 10% formalin for 30 minutes and washed twice with 2 mL of double-deionized water. Culture plates were exposed to acetic acid from the kit (4-5 drops) for 3 minutes; acid was removed by tipping the excess off. Alcian blue solution was added to the culture and left at room temperature for 35 minutes. Excess stain was removed by aspirating, and cells were rinsed in acetic acid for 20 seconds. Culture wells were rinsed 5 times with tap water (3 mL, removed by aspiration) and then rinsed once with 2 mL of double-deionized water. Fixed cultures were dehydrated with 2 ethanol washes (1 each with 70% and 100% ethanol). Excess ethanol was aspirated, and samples were air-dried for 15 minutes before imaging with light microscopy.

## Results

### Growth Kinetics of Human and Porcine ADMSCs

The confluence of hADMSCs and pADMSCs in passage 4 was compared as a function of time (Figure 2). A 1-way analysis of variance comparison showed no significant difference in growth between human cells cultured at 37°C and porcine cells cultured at 39°C when both were plated at an initial concentration of 20,000 cells/cm^2^ (95% CI, −13.20 to 9.65 hours). The time to reach 90% confluence at these conditions was 55 and 59 hours (∼2.5 days) for porcine and human cells, respectively. Porcine cells plated at high density and incubated at human body temperature of 37°C showed slower growth compared with human cells (95% CI, 25.81-48.93 hours) and porcine cells grown at 39°C (95% CI, 26.96-51.68 hours) (*P*<.001). At 37°C, the average time for porcine cells to reach 90% confluence was 176 hours (7.3 days).

**Figure 2.**
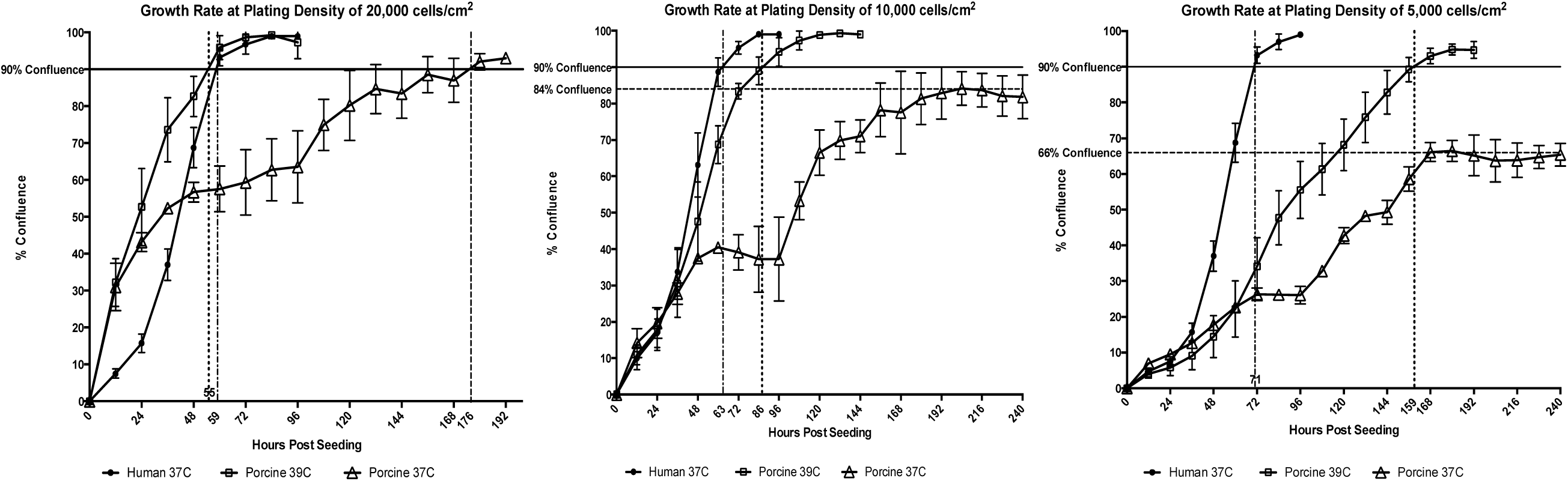
Growth Kinetics of Human and Porcine ADMSCs, Stratified by Plating Density. Cells were seeded at a high density (20,000 cells/cm^2^, left), medium density (10,000 cells/cm^2^, center), and low density (5,000 cells/cm^2^, right). After 72 hours, the porcine cells cultured at 39°C grew at a rate similar to that of human ADMSCs at 37°C when seeded at 20,000 cells/cm^2^ (*P*<.001; 95% CI, −13.20 to 9.65 hours). When cultured at 37°C or seeded at concentrations lower than 20,000 cells/cm^2^, porcine ADMSCs grew more slowly than human ADMSCs. Porcine ADMSCs did not reach 90% confluence when seeded at an initial concentration of 5,000 cells/cm^2^, even after 10 days of culture. ADMSC indicates adipose-derived mesenchymal stem cells.

When plated at medium or low initial density, porcine cells incubated at 39°C grew more slowly than hADMSCs (95% CI for medium density, 5.16-18.77 hours; low density, 48.66-69.55 hours). The average time to reach 90% confluence for hADMSCs and pADMSCs plated at medium density and incubated at the species-specific body temperatures was 63 and 86 hours, respectively.

In all cases, pADMSCs incubated at 37°C showed slower growth compared with hADMSCs incubated at 37°C and pADMSCs incubated at 39°C (Figure 2). Although pADMSCs initially plated at 20,000 cells/cm^2^ did reach 90% confluence after 7.3 days, those plated at lower densities never reached 90% confluence, even with an extended incubation of up to 10 days. The 37°C pADMSC group also showed visual signs of early detachment.

In addition to determining the overall growth curve, we also calculated the doubling time for hADMSCs and pADMSCs from the linear portion of the growth curve. Doubling times and linear regression values for each temperature and plating concentration are shown in Table 2. Cellular doubling time for pADMSCs improved with higher-temperature incubation by 74.3 to 162.2 hours, depending on initial plating density (*P*<.001).

**Table 2.** Doubling Times for Porcine and Human Adipose-Derived Stem Cells Cultured in 5% Human Platelet Lysate−Enhanced Media

| Cell Type and<br>Growth<br>Temperature | Seeding Concentration, cells/cm<sup>2</sup> |  |  |  |  |  |
| --- | --- | --- | --- | --- | --- | --- |
|  | 20,000 |  | 10,000 |  | 5,000 |  |
|  | Doubling |  | Doubling |  | Doubling |  |
|  | Time, h | R<sup>2</sup> | Time, h | R<sup>2</sup> | Time, h | R<sup>2</sup> |
| Human 37°C | 11.1 | 0.997 | 14.6 | 0.989 | 11.1 | 0.996 |
| Porcine 39°C | 20.1 | 0.988 | 17.8 | 0.995 | 19.6 | 0.999 |
| Porcine 37°C | 173.3 | 0.947 | 88.9 | 0.781 | 110 | 0.819 |

### Trilineage Differentiation of Human and Porcine ADMSCs

Trilineage differentiation of human and porcine MSCs was performed by using StemPro differentiation media. Human samples incubated at 37°C showed positive staining for adipogenesis and osteogenesis after 14 days in differentiation media (Figure 3). We encountered difficulties in obtaining positive trilineage differentiation when incubating porcine cells at 37°C. Most samples showed signs of early detachment (starting on day 7), although we observed very weak osteogenic differentiation on day 14. None of our osteogenic cultures remained attached beyond day 15, despite using Cell-Bind plates (Corning Scientific; Corning, NY) to improve attachment efficiency. All micromass cultures for porcine chondrogenesis failed because of detachment by day 14 of incubation at human body temperature.

**Figure 3.**
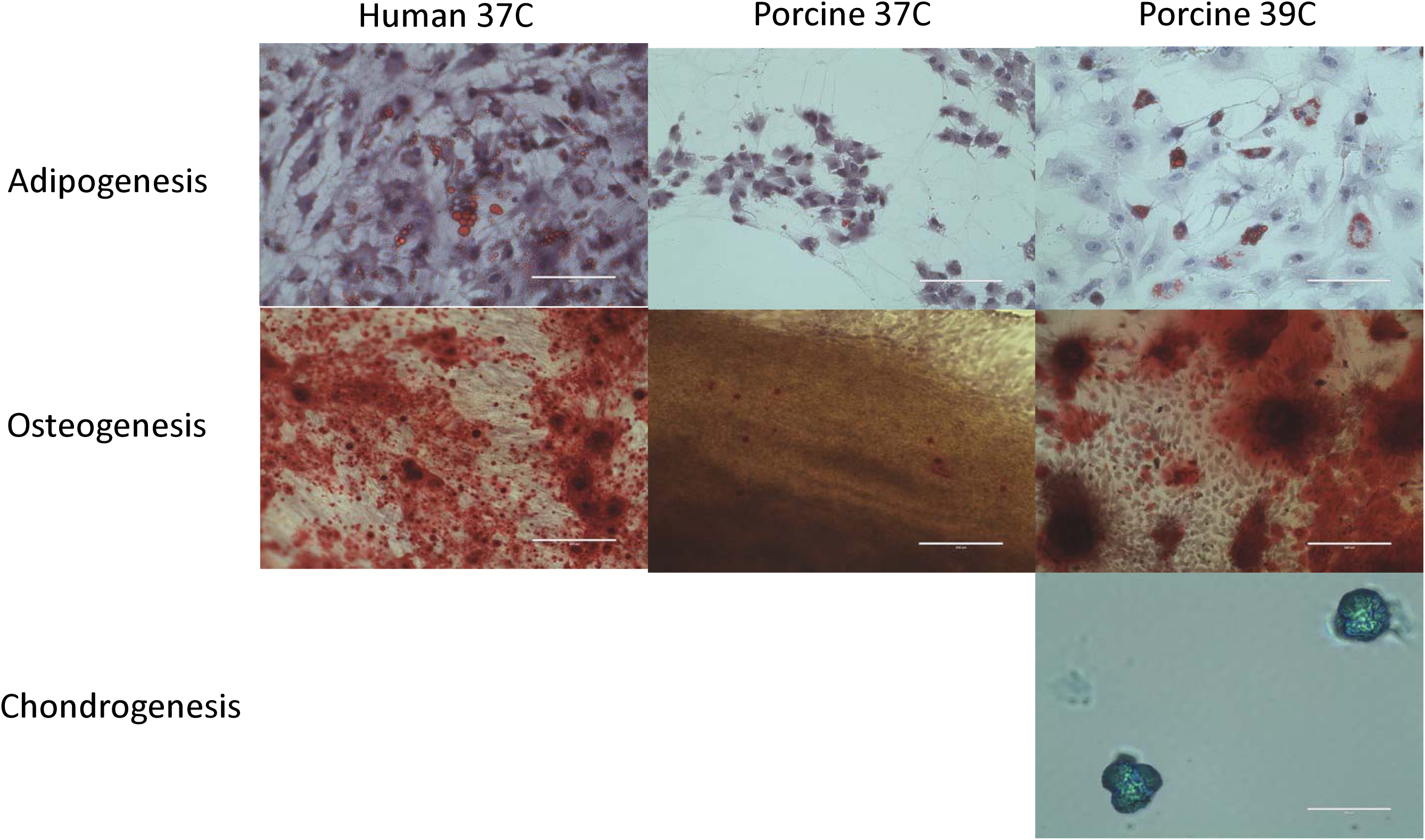
Trilineage Differentiation of Porcine Adipose−Derived Mesenchymal Stem Cells at 39°C vs 37°C. Standard stains for trilineage differentiation were used: Oil-Red-O for adipocytes, Alazarin red for osteocytes, and Alcian blue for chondrocytes. At 37°C, porcine cells had very poor adherence to the culture dish, and differentiation into adipocytes or osteocytes was questionable. Chondrocyte micromass cultures detached from culture plates under all 37°C conditions and did not undergo complete differentiation. At 39°C, the cells’ ability to differentiate into fat, bone, and cartilage markedly improved.

The higher incubation temperature of 39°C showed significant improvement in adipogenesis, osteogenesis, and chondrogenesis for pADMSCs. Strong adipogenesis and osteogenesis was observed on days 14 and 18, respectively. Osteogenic cultures were positive on day 14, albeit not as strongly differentiated as they were on day 18 (data not shown). Chondrogenic cultures were allowed to grow for 21 days and showed positive staining with Alcian blue in all cultures (Figure 3).

## Discussion

We described how to harvest fat from either the back or chest of domestic swine. From our surgical experience, we recommend harvesting from the back for most pigs. If the back will be used for testing and cannot be disturbed, the upper chest is an alternative area for adipose harvest. Unlike humans, domestic swine have much smaller fat reserves in the abdomen when compared with the back or the chest [16].

The growth characteristics and trilineage differentiation data show that incubation at 39°C is necessary for optimal in vitro cellular performance. Our confluence data at varying temperatures and our analysis of doubling times show that pADMSCs function better when incubated at normal porcine body temperature (39°C) compared with human body temperature (37°C). At the higher temperature, the doubling time of pADMSCs decreased by nearly one-third compared with growth at 37°C. Likewise, incubation close to the normal body temperature markedly improved the cells’ ability to differentiate into fat, bone, and cartilage.

Unlike human cells, which seemed to multiply almost equally well at high-, medium-, and low-density plating concentrations, porcine cells showed better growth at medium to high plating densities. This effect is likely due to a required paracrine effect. We used a human platelet lysate in the porcine culture media, at the same concentration used to culture human cells. It could be that this density requirement could be overcome by using higher concentration of platelet lysate or by using pig platelet lysate. The need for a higher initial plating concentration for growth translates into requiring more harvested fat compared with human donors or ensuring that cells are maintained at a higher concentration after plating. We recommend harvesting at least 7.5 g and optimally at least 10 g to seed 2 T-75 flasks at passage zero.

### Conclusion and Final Recommendations

We believe that in vitro and in vivo experiments will have different or improved outcomes if porcine cells are incubated at 39°C instead of 37°C. Taken together, these data show that pADMSCs require higher initial plating densities and incubation at 39°C for the reproducible and predictable use of an autologous large-animal model for testing ADMSC therapies.

## Abbreviations

ADMSC: adipose-derived mesenchymal stem cell
hADMSC: human adipose−derived mesenchymal stem cell
MSC: mesenchymal stem cell
pADMSC: porcine adipose−derived mesenchymal stem cell
PBS: phosphate-buffered saline

